# Dietary Resilience of Termite Gut Microbiota and Enzymatic Function Reflects Feeding Strategy

**DOI:** 10.1101/2025.05.07.651711

**Authors:** Letícia Menezes, João Paulo L. F. Cairo, Ana Maria Costa-Leonardo, Maria Teresa Pedrosa Silva Clerici, Isabela da Costa Barreto, Bianca Suriano Francisco dos Santos, Alberto Arab

## Abstract

Termites are major decomposers in tropical ecosystems, relying on complex gut microbiomes to digest lignocellulosic substrates. In this study, we compared the gut microbiota composition and enzymatic responses to dietary shifts in two neotropical termite species with contrasting feeding strategies: the polyphagous *Silvestritermes euamignathus* and the litter-feeding specialist *Cornitermes cumulans*. High-throughput sequencing and enzymatic assays revealed that *S. euamignathus* maintained stable microbial communities and enzymatic profiles across diverse diets, including artificial and fiber-rich substrates. In contrast, *C. cumulans* exhibited significant shifts in bacterial abundance and reduced enzymatic activity under altered diets, particularly those differing from its natural litter-based diet. Functional gene predictions further indicated broader metabolic potential in *S. euamignathus*, particularly in response to complex substrates, while *C. cumulans* showed transcriptional suppression of polysaccharide-degrading enzymes. These results suggest that *S. euamignathus* benefits from a more flexible and functionally resilient gut symbiosis, enabling adaptation to heterogeneous or disturbed environments. In contrast, the narrower metabolic scope of *C. cumulans* may limit its capacity to respond to dietary change. Our findings highlight how feeding ecology shapes microbiome plasticity and digestive function in termites, with implications for understanding their adaptability under environmental stress and climate-driven shifts in resource availability.

## 1 Introduction

Termites are key decomposers in tropical ecosystems, relying on complex gut microbiomes to digest lignocellulosic materials. While non-termitid species primarily consume wood and depend on a combination of protists, bacteria, and archaea for digestion, termitids (Termitidae) rely almost exclusively on a highly diverse bacterial community. This microbial diversification has been closely linked to the evolutionary success of termitids, enabling them to exploit a wide range of ecological niches and feeding substrates.(Brune, 2014; Chouvenc et al., 2021; Arab et al., 2024).

Feeding habits among termitids vary considerably, ranging from dietary specialists that primarily consume litter, grass, or soil to polyphagous generalists capable of accessing a broader range of lignocellulosic resources (Mikaelyan et al., 2015; Moreira et al., 2018). This trophic diversity likely co-evolved with distinct microbial assemblages and enzymatic capabilities across termite lineages. Generalist feeders may harbor more plastic and resilient microbiomes, providing metabolic flexibility under fluctuating environmental conditions. In contrast, specialist feeders may exhibit more rigid symbiotic associations, shaped by long-term dietary specialization (Macke et al., 2017). These ecological contrasts offer an ideal framework for testing how feeding strategy in termites influences gut microbiota composition and digestive function.

Microbiota plasticity is recognized as a critical factor in insect’s ability to cope with dietary shifts (Engel and Moran, 2013). In termites, previous studies suggest that changes in diet can lead to rapid, but often reversible, shifts in gut microbial communities (Miyata et al., 2007; Wang et al., 2016). However, most of this research has focused on a limited number of species, and little is known about the flexibility of gut microbiota and associated enzymatic traits in neotropical termites with contrasting diets.

Here, we investigate how gut microbiota structure and enzymatic function respond to dietary manipulation in two neotropical termitids: *Silvestritermes euamignathus*, a polyphagous species with a broad feeding repertoire, including litter, wood, and the carton nest of other termites (Mathews, 1977), and *Cornitermes cumulans*, a litter specialist (Coles De Negret and Redford, 1982)(Figures 1A-B). We hypothesize that (1) *S. euamignathus* will exhibit greater stability in both microbial composition and enzymatic activity across diets, reflecting its dietary flexibility, while (2) *C. cumulans* will show significant microbial shifts affecting its functional profile when subjected to different diets. By comparing microbiota diversity, taxonomic responses, and enzymatic profiles across dietary treatments, we aim to better understand the ecological and evolutionary mechanisms related to digestive resilience in termites.

**Figure 1.**
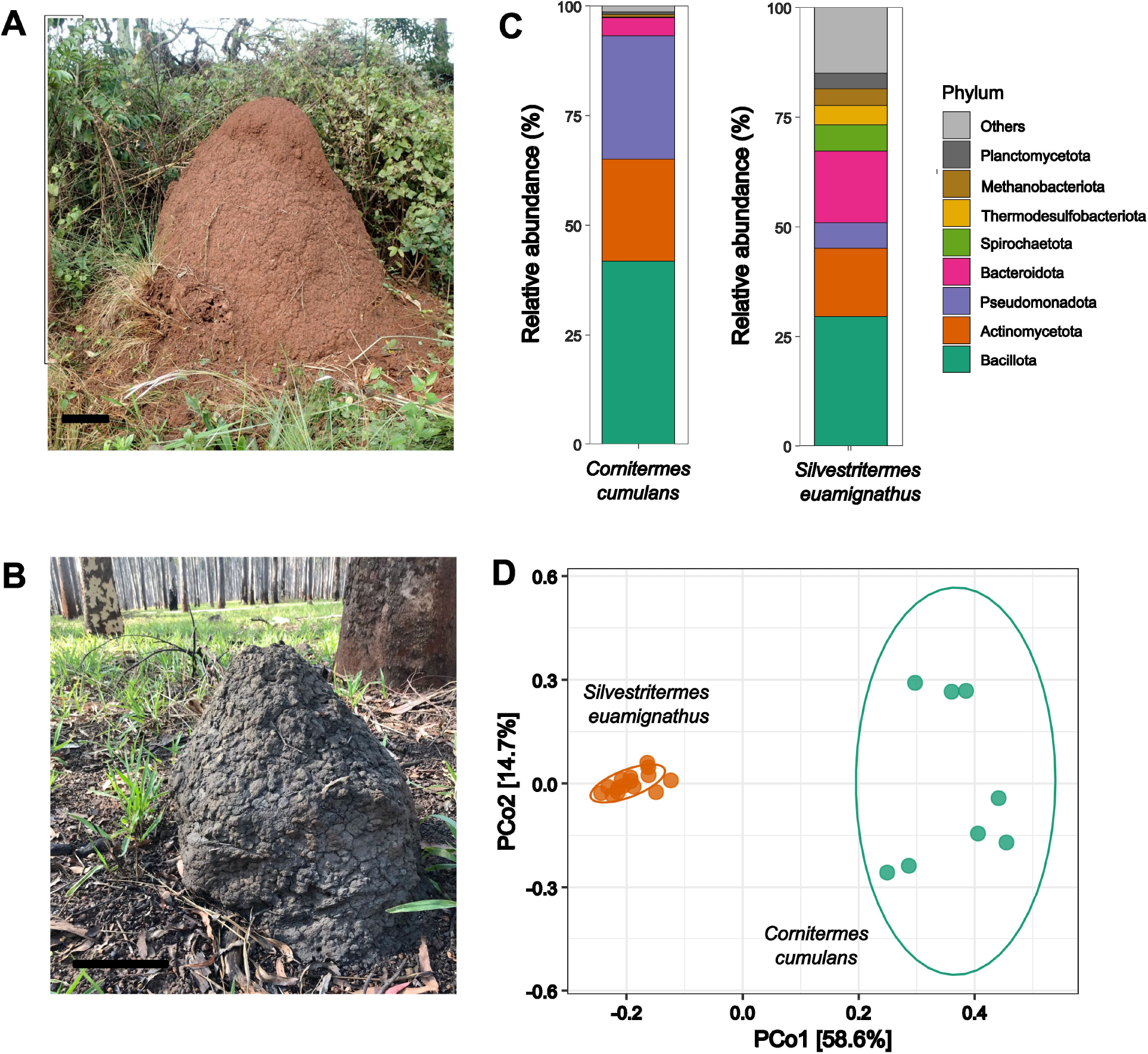
Epigeal nests of (A) *Silvestritermes euamignathus* and (B) *Cornitermes cumulans*. (C) Relative abundance of major bacterial phyla in the worker gut microbiota. (D) Ordination plot showing differences of gut bacterial composition between the two termite species.

## 2 Materials and methods

### 2.1 Termites and study site

Colonies of *Silvestritermes euamignathus* and *Cornitermes cumulans* (Termitidae: Syntermitinae) were collected from pasture areas in the Campinas-SP region (22° 46’29.0”S, 47° 04’06.5”W). The nests were kept in 50 L plastic containers in a climate-controlled room at 26°C, under a 12-hour photoperiod cycle, with water and grass provided as food. Sampling was authorized by the Brazilian Institute of Environment and Renewable Natural Resources (IBAMA), the enforcement agency of the Brazilian Ministry of the Environment (SISBIO permit no. 72676-4).

### 2.2 Feeding experiments

Feeding experiments were performed using artificial diets differing in carbohydrate complexity. From each of four colonies, 50 workers and five soldiers were carefully extracted and placed in 10 cm Petri dishes containing sterilized, moistened sand and a 3 cm potato dextrose agar disc supplemented with one of the following treatments:

Dextrose (40% dextrose + 15% agar), a monosaccharide representing a simple carbohydrate.

CreaFibe (40% CreaFibe QC 90 + 15% agar), comprising 84–88% cellulose, 12–16% hemicellulose, and <1% lignin (manufacturer’s specifications).

Arbocel® RC fine lignocellulose (40% Arbocel + 15% agar), containing 65–70% insoluble cellulose and >20% lignin (Hussein et al., 2017).

Control (15% agar only).

After five days, surviving termites were enumerated, and their digestive tracts were dissected and stored at −22°C for enzymatic and molecular analyses (Menezes et al., 2018). An in-situ sample (50 workers per colony) was retained without experimental manipulation. Food consumption was quantified as the difference between initial and final fresh weight of diet discs, adjusted for humidity loss using control dishes (food without termites). Differences in consumption and survival among treatments were assessed via Generalized Linear Mixed Models (GLMMs) with Gaussian (consumption) and Poisson (survival) error distributions, implemented in the glmmTMB package (Brooks, Mollie et al., 2017). Pairwise comparisons were conducted using estimated marginal means with th emmeans package (Lenth et al., 2023), and residual distributions were evaluated with the DHARMa package (Hartig, 2022).

### 2.3 DNA extraction

The entire guts of 50 workers from each of the three colonies subjected to each food treatment and from the *in situ* sample were placed in 2 mL tubes containing 1 mL of lysis buffer (500 mM NaCl, 50 mM Tris-HCl, pH 8.0, 50 mM EDTA, and 4% sodium dodecyl sulfate). DNA extraction was performed using a bead-beating protocol as previously described [27]. DNA integrity was assessed via agarose gel electrophoresis (1.0% w/v), and quantification was conducted using a NanoDrop spectrophotometer by measuring absorbance at 260 nm.

### 2.4 16S rRNA library preparation, sequencing, and taxonomic assignment

The V4 region of the 16S rRNA gene was amplified using primers 515F (5′ TCGTCGGCAGCGTCAGATGTGTATAAGAGACAGGTGCCAGCMGCCGCGGTAA 3′) and 806R (5′ GTCTCGTGGGCTCGGAGATGTGTATAAGAGACAGGGACTACHVGGGTWTCTAAT 3′), with Illumina overhangs (underlined) (Menezes et al., 2018). Library preparation involved two PCR steps. In the first step, specific primers were used for each library with Phusion Polymerase (Thermo Scientific) and 40 ng of template DNA per sample. The amplification conditions were: initial denaturation at 98°C for 2 min, followed by 30 cycles of 98°C (30 s), 60.1°C (30 s), and 72°C (40 s), concluding with a final extension at 72°C for 5 min. The second PCR step incorporated Illumina sequencing adapters and dual index barcodes (Nextera Index Kit, Illumina) into the amplified libraries. This reaction used Phusion polymerase (Thermo Scientific) with 100 ng of purified PCR products from the first step as the template, along with indexing primers from Illumina. The amplification conditions included an initial denaturation at 98°C for 3 min, followed by 5 cycles of 98°C (30 s), 55°C (30 s), and 72°C (30 s), ending with a final extension at 72°C for 5 min. Each sample was amplified in triplicate. Pooled samples were purified using Agencourt AMPure Magnetic Beads (Beckman Coulter) and quantified with a Qubit Fluorometer 2.0 (Invitrogen). Sequencing was performed on the Illumina MiSeq platform at the Brazilian Bioethanol Science and Technology Laboratory (CTBE/CNPEM) using a MiSeq Reagent Kit V3 (600 cycles). The dataset was processed using the DADA2 1.16 pipeline (Callahan, 2021). Taxonomic assignment was performed using the SILVA reference database (Quast et al., 2013). Tables generated by DADA2 pipeline were imported into phyloseq (McMurdie and Holmes, 2013).

### 2.6 Microbial diversity and community structure analyses

We used R version 3.5.0 (R Core Team, 2021) to conduct analyses using different packages. Downstream analysis, including α- and β-diversity analysis was calculated using the microeco and MicrobiotaProcess packages (Liu et al., 2021; Xu et al., 2023). The ASVs table was merged with relevant metadata into a microeco object. Generalized linear mixed models (GLMM) were performed to check for overall significant differences of α-diversity estimates and the total number of ASVs among samples. We fitted multivariate generalized linear models (mvabund package) (Wang et al., 2012) to test the effects of the diet on the microbial relative abundance. Models were fitted with a negative binomial distribution using 999 bootstrap iterations. The manyglm function was used to carry out the analysis. Ordination analyses were used to evaluate microbial composition at the ASV level. We used GLMM analysis to identify ASVs that define the differences between microbiomes. The potential genes encoding carbohydrate active enzymes (CAZy) were inferred using the Tax4Fun2 version 1.1.6 package (Aßhauer et al., 2015) and the reference profiles (Ref99NR) from the Kyoto Encyclopedia of Genes and Genomes (KEGG) database. Similarities greater than 97% were considered orthologous KEGG groups (KOs). The output included individual pathways (three levels) and related CAZy encoded genes (https://www.cazy.org/).

### 2.7 Assays of lignocellulolytic activity of termite guts

The enzymatic activity against cellulose and hemicellulose substrates was evaluated using crude enzyme extracts from the gut of workers from four colonies, each subjected to different dietary treatments, following the methodology described by Franco Cairo et al. (2011). The assays aimed to assess the activity of the soluble fraction of protein extracts against both natural polysaccharides and synthetic oligosaccharides with varying monomer compositions and branching patterns. Crude protein extraction was performed using the guts of 50 workers from each colony. The samples were homogenized in 2 mL of 100 mM sodium acetate buffer (pH 5.5) and then centrifuged at 20,100 × g for 30 minutes at 4°C. The supernatant was collected, and 1 µL of Protease Inhibitor Cocktail (Anresco) was added per mL of crude extract to prevent protein degradation. Protein concentration in each extract was determined using the Bradford method (Bradford, 1976).

Each enzymatic reaction consisted of 10 µL of crude protein extract, incubated at 37°C for 40 minutes with 40 µL of 50 mM sodium acetate buffer (pH 5.5) and 50 µL of a 0.5% specific carbohydrate solution (in water), performed in triplicate (Franco Cairo et al., 2011). To terminate the enzymatic reactions, 100 µL of dinitrosalicylic acid (DNSA) was added, followed by heating at 99°C for 5 minutes. The resulting color change was measured at 412 nm using a TECAN M2000 plate reader. Enzymatic activity was expressed in mmol of glucose equivalents produced per mg of protein. One unit of enzymatic activity was defined as the amount of enzyme required to release 1 mmol of reducing sugar. The carbohydrate substrates used in the enzymatic assays included CMC (carboxymethyl cellulose, low viscosity) (β-1,4-carboxymethylglucan), β-glucan from barley (β-1,4-glucan), xylan from oat spelt (β-1,4-xylan), rye arabinoxylan (α-2,3-arabinose-β-1,4-xylan) from citrus, lichenin from moss, and pNP-G (4-nitrophenyl β-1,4-D-glucopyranoside). Blank reactions were prepared using the same procedure, except that DNSA was added before incubation with the protein extract. The color change was analyzed at 412 nm using a TECAN M2000 plate reader. Glucose and p-nitrophenyl were used for standard curve construction. To evaluate the effect of diet on the enzymatic activity of termite workers, a Wilks’ Lambda-type nonparametric multivariate inference was conducted using the npmv package (Ellis et al., 2017).

## 3 Results

### 3.1 The gut microbiota of *Silvestritermes euamignathus* and *Cornitermes cumulans*

We identified a total of 2,222 bacterial ASVs (508,218 reads) and 90 archaeal ASVs (18,147 reads) in the gut microbiota of *S. euamignathus* workers, as well as 982 bacterial ASVs (289,091 reads) and 52 archaeal ASVs (14,812 reads) in *C. cumulans* workers (Tables S1 and S2). The most abundant phyla—Bacillota, Actinomycetota, and Bacteroidota—accounted for 67% and 69% of sequence reads in *S. euamignathus* and *C. cumulans*, respectively (Figure 1C). At the genus level, *Papillibacter* was the most abundant in *S. euamignathus*, while *Acinetobacter* dominated in *C. cumulans* (Figure S1A). The gut microbiota composition differed significantly between these termite species (*manyglm*; Dev = 35,21; df = 1; *p* < 0.001) (Figure 1D), with only 6.3% of ASVs shared between them (Figure S1B). Diversity indexes were not significantly different between the two termite species (Figure S1C and Table S3).

### 3.2 Diet Alters Bacterial Abundance in *C. cumulans* but Not in Polyphagous *S. euamignathus*

Diet variation had no significant effect on food consumption or survival in either the litter-feeding *Cornitermes cumulans* or the polyphagous *Silvestritermes euamignathus* (Figure S2 and Table S4). However, in *C. cumulans*, artificial diets led to notable shifts in the relative abundance of specific bacterial taxa, particularly within the phyla Bacillota and Pseudomonadota, which declined under artificial feeding regimes. In contrast, *S. euamignathus* showed no such marked variation in taxonomic abundance across diets (Figures 1A–B, S3; Tables S1 and S2). Despite these changes in bacterial abundance, gut microbiota composition remained unchanged in both species (*S. euamignathus*: Deviance = 9.292, p = 0.068; *C. cumulans*: Deviance = 4.890, p = 0.092) (Figures 2B–C). These results suggest that while *C. cumulans* exhibits shifts in specific microbial taxa in response to diet, its overall community structure remains relatively resilient, and *S. euamignathus* maintains both stable abundance and composition under dietary stress.

**Figure 2.**
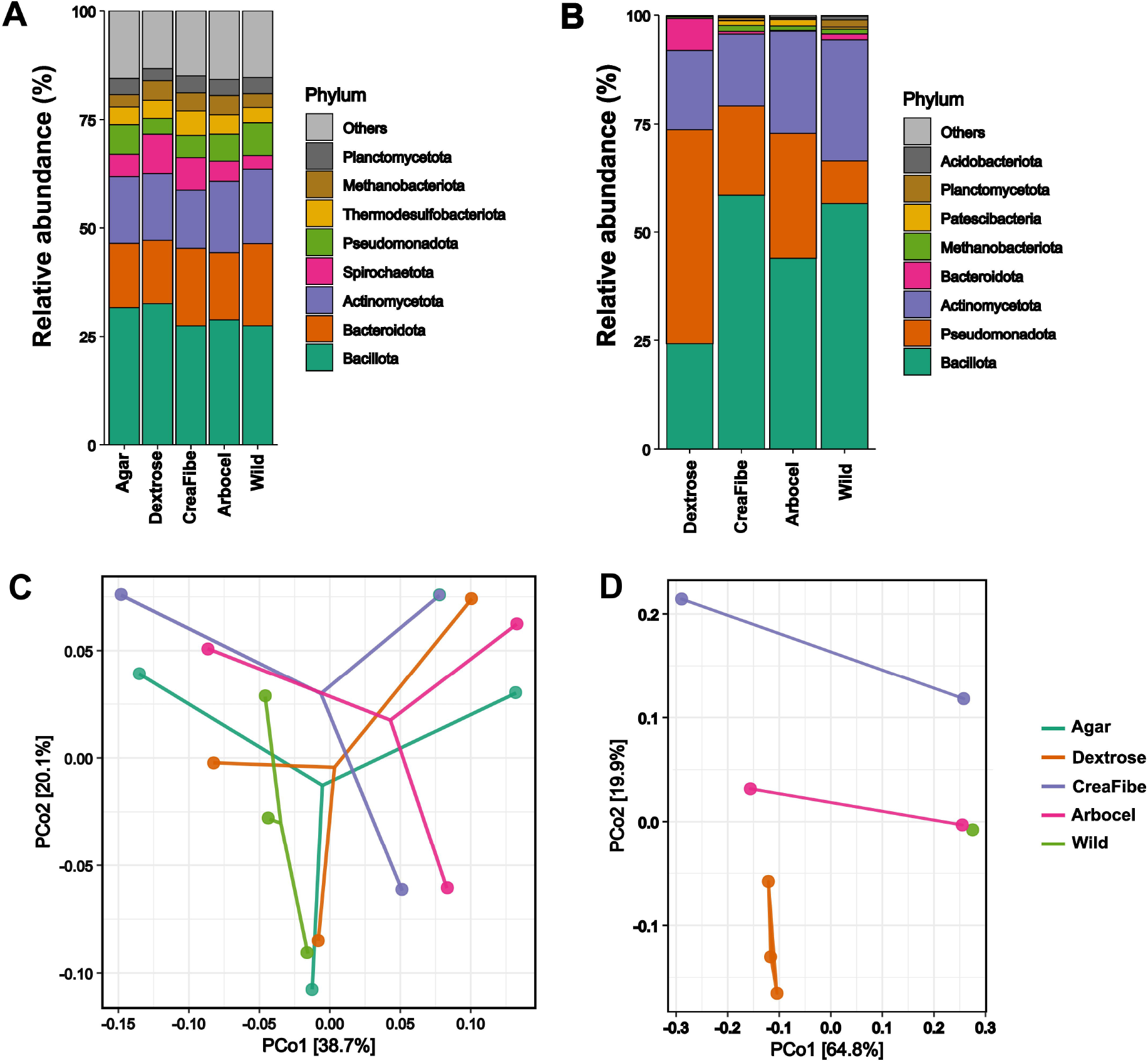
Effect of dietary treatments on the gut microbiota of *Silvestritermes euamignathus* and *Cornitermes cumulans*. Relative abundance of bacterial phyla in the gut microbiota of *S. euamignathus* (A) and *C. cumulans* (B) across different diets. Principal Coordinates Analysis (PCoA) based on weighted Unifrac distances showing beta diversity patterns for (C) *S. euamignathus* and (D) *C. cumulans*. Wild treatment consists in workers without experimental manipulation. We did not detect consumption of the control treatment (pure agar) by workers of *C. cumulans*.

### 3.3 Enzymatic Responses Reveal Greater Dietary Resilience in *S. euamignathus*

The functional inference of gut microbiota revealed differences between the two termite species in response to dietary treatments. In *S. euamignathus*, the dextrose diet strongly induced enzymes involved in simple sugar metabolism, including alpha-amylase (EC. 3.2.1.1) and alpha-glucosidase (EC. 3.2.1.20) (Figure 3A). However, individuals fed complex fiber-rich diets such as CreaFibe and Arbocel, or the wild treatment, showed moderate to higher profiles of ligninases (EC. 1.11.1.9 and EC. 1.11.1.15) and hemicellulases (EC. 3.2.1.10 and EC. 3.2.1.23) (Table S5). Enzymatic activity assays also showed consistent profiles across dietary treatments, although individuals fed the dextrose diet exhibited slightly elevated activity for some substrates, retaining higher activity on β-glucan and pNP-G substrates even when exposed to artificial diets (Wilks’ Lambda = 7.91; df = 15, 36.29; p < 0.001) (Figure 3C and Table S6).

**Figure 3.**
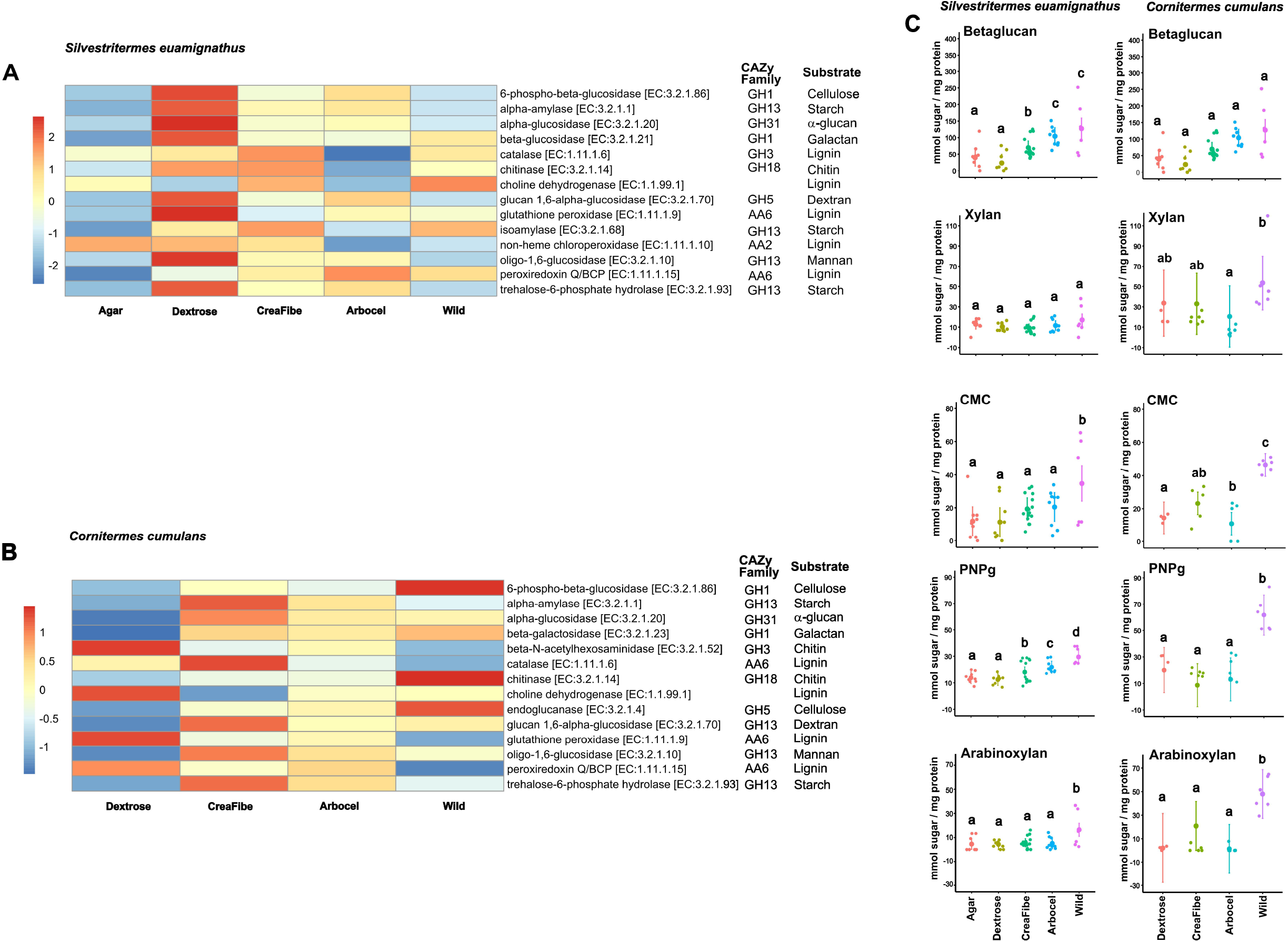
Heatmaps showing the expression of enzymes related to carbohydrate and lignin degradation predicted by Tax4Fun across dietary treatments for (A) *Silvestritermes euamignathus* and (B) *Cornitermes cumulans*. (C) Quantification of enzymatic activities (nmol sugar/mg protein) of the soluble fraction of gut protein extracts of each termite specie. Substrates abbreviations: CMC (carboxymethyl cellulose) (β-1,4-carboxymethylglucan); β-glucan (β-1,4-glucan), Xylan (β-1,4-xylan), Arabinoxylan (α−2,3-arabinose-β-1,4-xylan), and pNP-G (4-nitrophenyl – β-1,4-D-glucopyranoside). Different letters indicate significance at P< 0.05 (lsmeans pairwise analysis). Wild treatment consists in workers without experimental manipulation.

In contrast, *Cornitermes cumulans* displayed a pronounced reduction of enzymes involved in polysaccharide degradation—particularly those related to starch, cellulose, and hemicellulose—when exposed to artificial diets (Figure 3B and Table S7). This functional suppression was reflected in significantly lower enzymatic activities against substrates such as xylan, CMC, PNPg, and arabinoxylan compared to wild-fed individuals (Wilks’ Lambda = 3.14; df = 20, 130.29; p < 0.001) (Figure 3D and Table S8). Altogether, these results demonstrate that *S. euamignathus* is more resilient to dietary shifts, maintaining functional digestive enzyme activity across a range of nutritional environments. *C. cumulans*, in contrast, exhibits more pronounced reductions in enzymatic performance under simplified or artificial diets, suggesting a narrower metabolic scope and lower tolerance to dietary change.

## 4 Discussion

Our findings demonstrate that the polyphagous termite *Silvestritermes euamignathus* exhibits greater stability in both gut microbiota composition and enzymatic function across dietary treatments compared to the litter-feeding specialist *Cornitermes cumulans*. This supports the idea that microbial diversity enhances host resilience to nutritional variability (Bertino-Grimaldi et al., 2013; Macke et al., 2017). The stability observed in *S. euamignathus* mirrors patterns in other generalist insects, where high microbial diversity provides metabolic flexibility under novel or suboptimal diets (Shukla et al., 2016; Tinker and Ottesen, 2016). In contrast, *C. cumulans* exhibited significant shifts in microbial abundance and a marked decline in enzymatic activity when exposed to artificial diets. These findings suggest a symbiotic system adapted to a narrower dietary niche, consistent with its evolutionary specialization on litter.

Our results build upon previous studies showing that termite gut microbiomes are shaped by both host phylogeny and feeding ecology (Mikaelyan et al., 2015). In wood-feeding termites, for instance, dietary changes often lead to transient microbial shifts that revert after reestablishing the native diet (Miyata et al., 2007; Wang et al., 2016). We expand this view by showing that *S. euamignathus*, a generalist feeder, maintains a stable microbiota and consistent enzyme profiles remain largely stable across a range of dietary conditions, suggesting a more flexible symbiotic system across diverse substrates, suggesting a more flexible and functionally redundant symbiotic system. Meanwhile, *C. cumulans* exhibits reduced microbial diversity and enzymatic function under dietary change, consistent with the constrained plasticity observed in other specialists (Colman et al., 2012; Pérez-Cobas et al., 2015; Macke et al., 2017).

Importantly, our data reveal that dietary complexity influences not only microbial composition but also functional gene predictions and host enzymatic responses. In *S. euamignathus*, complex diets (e.g., CreaFibe, Arbocel, and the wild substrate) triggered broader CAZy functional profiles and elevated expression of lignocellulolytic enzymes. These responses likely reflect adaptation to a naturally heterogeneous diet and the capacity to regulate microbial function according to substrate complexity (presence of hemicellulose and lignin). By contrast, *C. cumulans* showed functional suppression and reduced enzyme activity when fed artificial fiber-rich diets—indicating limited capacity to upregulate digestion in response to unfamiliar or compositionally complex substrates.

These findings highlight that resilience to dietary shifts is not only a matter of microbial diversity, but also of functional plasticity—the ability to activate or suppress microbial and enzymatic pathways in response to nutritional cues (Boucias et al., 2013). This adaptive flexibility may contribute to the ecological success of generalists like *S. euamignathus* in disturbed or changing environments.

Finally, these patterns have broader ecological and evolutionary implications. In the context of global climate change and increasing anthropogenic disturbance, termites with flexible gut microbiomes may better withstand shifts in resource availability. As plant biomass becomes more accessible under warming scenarios, generalist species may expand into new habitats, facilitated by symbiotic versatility and metabolic robustness. Conversely, specialists like *C. cumulans* may face greater risk if their preferred substrates decline, potentially altering decomposition dynamics and species distributions (Gilbert et al., 2015; Buczkowski and Bertelsmeier, 2017). Understanding how diet shapes gut symbiosis is thus crucial for predicting termite responses to ecological change.

## 6 Author Contributions

The authors AA and AMCL designed the study. LM, JPFC, MTPSC, IDCB, BSFS, and AA performed the experiments. AA analysed data. AA and AMCL wrote the paper.

## 8 Acknowledgments

We would like to thank Brazilian Biorenewables National Laboratory (LNBR/CNPEM) NGS Sequencing Facility for generating the sequencing data described here. We are also grateful to Nutrassim Food Ingredients for providing the CreaFibe used in the dietary treatments.

## 10 Figure Captions

**Figure S1**. (A) Relative abundance of bacterial genera in the gut microbiota of *S. euamignathus* and *C. cumulans* across different diets. (B) Venn diagram of Amplicon Sequence Variants (ASVs) between the two termite species. (C) Alpha diversity indexes of gut microbiota between the two termite species.

**Figure S2**. Consumption and survival of workers of *S. euamignathus* and *C. cumulans* across different diets.

**Figure S3**. Alpha diversity indexes of *S. euamignathus* and *C. cumulans* worker gut microbiota across different diets.

## 11 Data Availability Statement

The datasets generated for this study can be found in the supplementary section. Termite metagenome sequences are deposited at https://www.ncbi.nlm.nih.gov/sra/PRJNA1256848

