## Supplementary figures and images for "Dietary Resilience of Termite Gut Microbiota and Enzymatic Function Reflects Feeding Strategy"

### Supplemental Figures

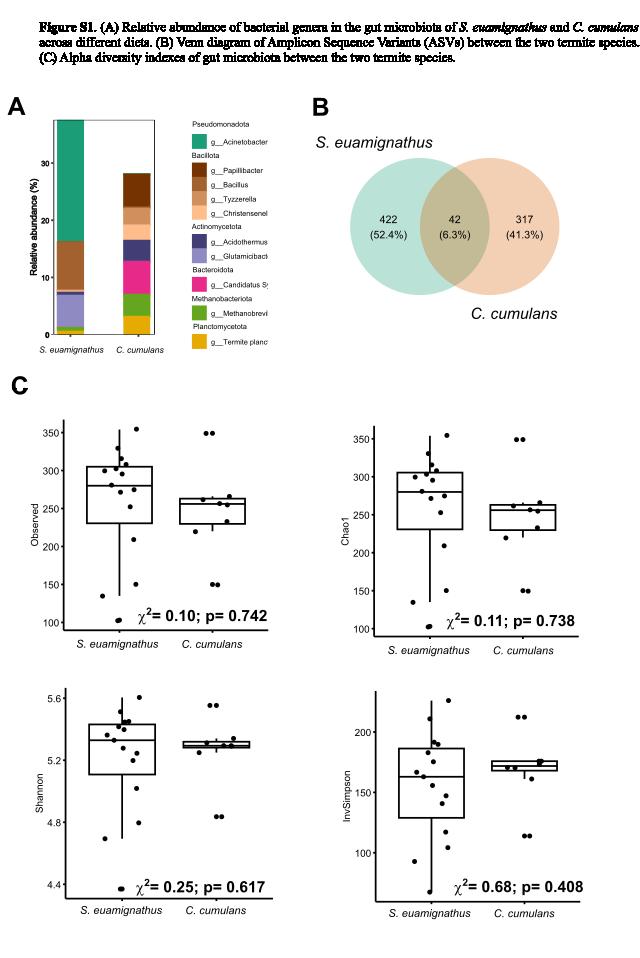

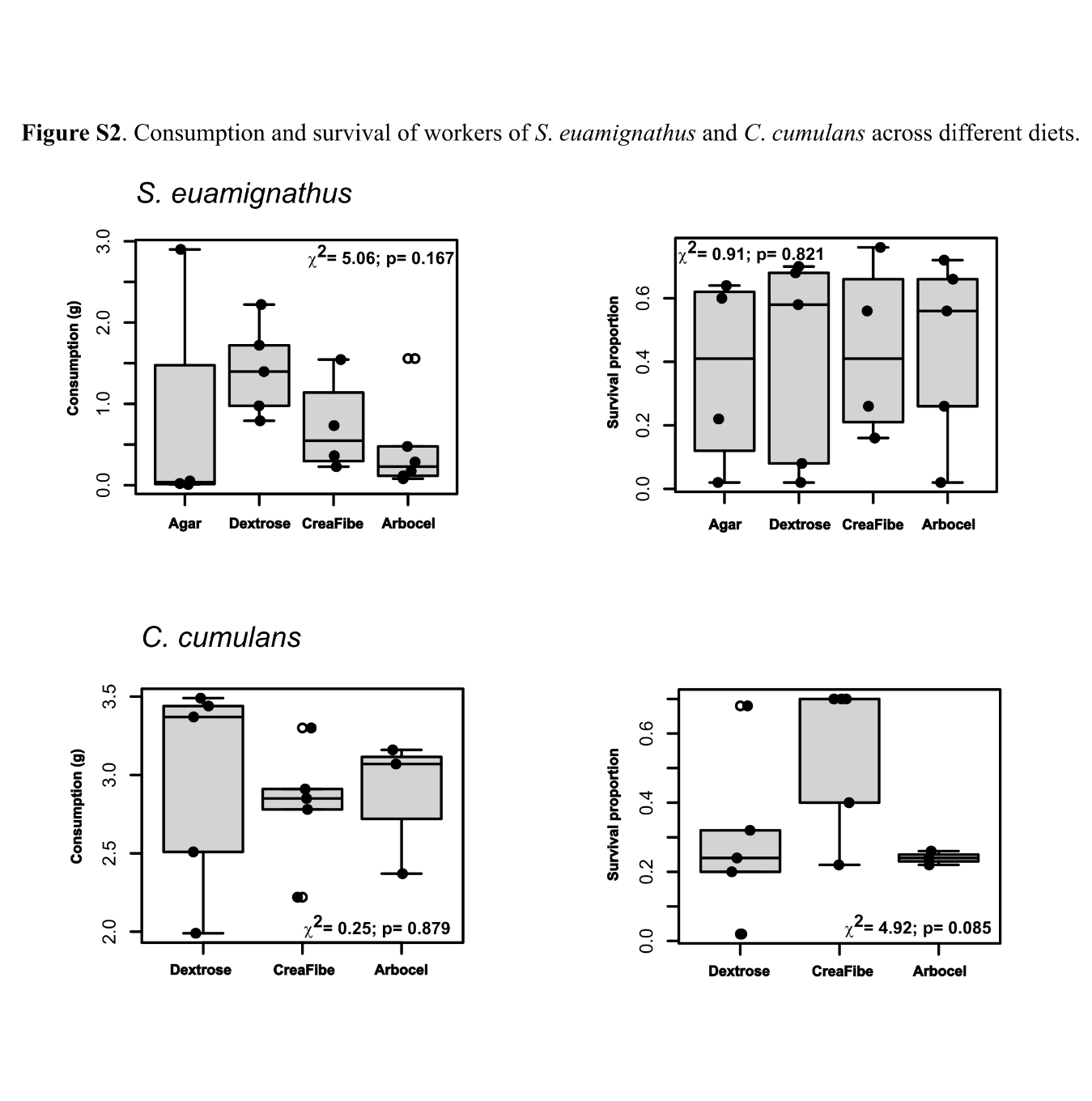

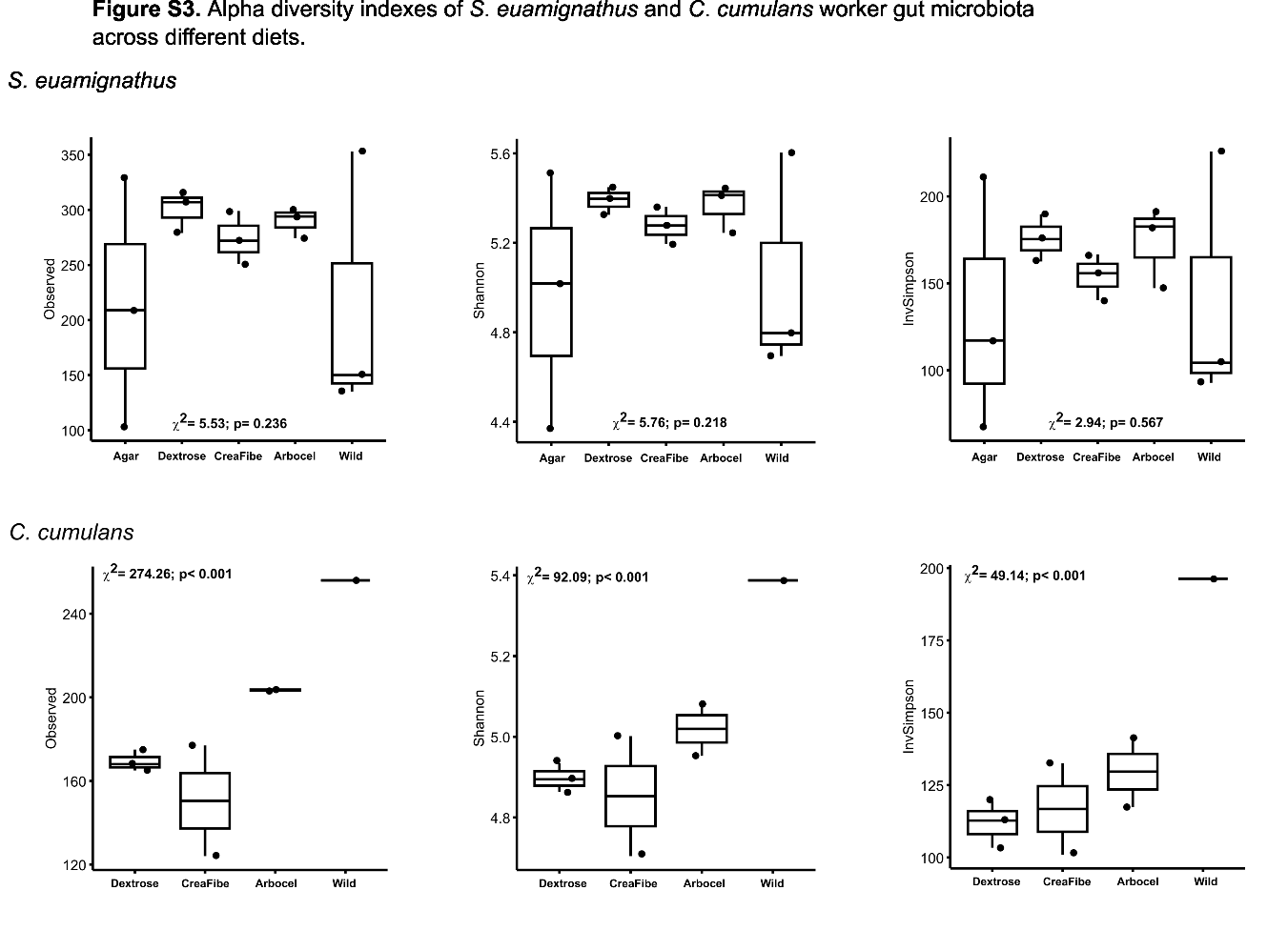
